# Endocytosis frustration potentiates compression-induced receptor signaling

**DOI:** 10.1101/2019.12.19.883066

**Authors:** Francesco Baschieri, Dahiana Le Devedec, Nadia Elkhatib, Guillaume Montagnac

**Author notes:** These authors contributed equally (Francesco Baschieri, Dahiana Le Devedec).

## Abstract

Cells experience mechanical stresses in different physiological and pathological settings. Clathrin-coated structures (CCSs) are sensitive to such perturbations in a way that often results in a mechanical impairment of their capacity to bud, ultimately impairing endocytosis. Compressive stress is a particular mechanical perturbation that leads to increased membrane tension and promotes proliferative signals. Here, we report that compression leads to CCSs frustration and that CCSs are required to potentiate receptor-mediated signaling in these conditions. We first confirmed that pressure stalls CCSs dynamics and showed that it also slows down the dynamic exchange of CCSs building blocks. As previously reported, compression-induced paracrine activation of the epidermal growth factor receptor (EGFR) was the primary cause of ERK activation in these conditions. We observed that the EGFR was efficiently recruited at CCSs upon compression and that CCSs were required for full ERK activation. In addition, we demonstrated that compression-induced frustrated CCSs could also serve as signaling platforms for the hepatocyte growth factor receptor (HGFR), provided HGF was present in the medium. We thus propose that, besides the particular case of EGFR paracrine activation, CCS frustration resulting from mechanical perturbations can potentiate signaling through different receptors with potential important consequences on cell adaptation to its environment.

## INTRODUCTION

Clathrin-mediated endocytosis (CME) relies on the assembly of clathrin-coated structures (CCSs) at the internal leaflet of the plasma membrane. CCSs are endowed with the capacity to recruit specific receptors and to bend the membrane in order to generate receptor-containing endocytic vesicles (McMahon and Boucrot, 2011). Membrane bending is however sensitive to mechanical perturbations that oppose the invagination force generated by CCSs. For instance, high plasma membrane tension stalls CCSs invagination and thus prevents CME (Boulant et al., 2011; Raucher and Sheetz, 1999). Other types of mechanical perturbations can also prevent normal CCSs budding. For example, a subset of CCSs termed tubular clathrin/AP-2 lattices (TCALs) that specifically nucleate at cell/collagen fibers contact sites show reduced dynamics because they try and fail to internalize fibers that are longer than the cell itself (Elkhatib et al., 2017). High substrate rigidity can also impair CCSs budding through favoring the αvβ5 integrin-dependent formation of flat and long-lived clathrin-coated plaques (Baschieri et al., 2018). Thus, CCSs frustration is a common response to a wide array of mechanical perturbations. CCSs frustration may not simply be a passive consequence of environmental perturbations but may actually participate in building an adapted response to these modifications. Indeed, we showed that TCALs help the cell to migrate in 3D environments and that clathrin-coated plaques that assemble on stiff substrates serve as signaling platforms for different receptors, ultimately leading to sustained cell proliferation (Baschieri et al., 2018; Elkhatib et al., 2017).

Cell compression was recently shown to induce CCSs frustration as well, most likely because of an increased membrane tension (Ferguson et al., 2017). Compressive forces are frequently encountered in the organism, whether in a physiological or pathological context (Butcher et al., 2009; Kalli and Stylianopoulos, 2018; Nia et al., 2017; Rakesh et al., 2010; Tschumperlin et al., 2004). These forces deeply impact the cell physiology and modulate signaling pathways as well as gene expression profile. For example, in bronchial epithelial cells, compression was shown to activate the epidermal growth factor receptor (EGFR) through a force-induced, metalloprotease-dependent shedding of HB-EGF precursor (Tschumperlin et al., 2004). In addition, mechanical compression in solid tumors was reported to promote both proliferation and apoptosis via mechanisms that are still not entirely clear (Alessandri et al., 2013; Cheng et al., 2009). Yet, whether compression-induced EGFR activation (Tschumperlin et al., 2002; Tschumperlin et al., 2004) is at stake in tumors is not known. Because we previously observed that the EGFR uses frustrated clathrin-coated plaques as signaling platforms (Baschieri et al., 2018), we wondered whether pressure-induced CCSs frustration could participate in EGFR signaling in these conditions.

## RESULTS

### Compression reduces CCS dynamics

To investigate the consequences of compressive forces on CCSs dynamics, we used HeLa cells that were genome-edited to express a fluorescently-tagged version of µ2-adaptin, a subunit of the clathrin adaptor AP-2. These cells were grown on glass coverslips and confined under an agarose plug under a constant compressive stress. While compressive stress is negligible in healthy tissues (Nia et al., 2017), it ranges from 0.21 to 20 kilo Pascal (kPa) in different solid tumors (Fernández-Sánchez et al., 2015; Kalli and Stylianopoulos, 2018; Nia et al., 2017). Additionally, compression stress higher than 3.87 kPa were reported to increase apoptosis in tumor spheroids in vitro (Cheng et al., 2009). For the present study, we decided to use a constant solid stress of 1 kPa in all our assays. We noticed that confinement induced an enlargement of the cell area and blebs were often observed at the cell edges (Fig. S1a), suggesting that membrane tension is most likely dramatically increased in these conditions (Gauthier et al., 2012). In addition, the nuclei of cells under compression were enlarged and nuclear blebs were also visible at their rim (Fig. S1b). These observations indicate that cells are indeed experiencing compression in our assays. In classical culture conditions, HeLa cells display a mixture of canonical, dynamic CCSs and static clathrin-coated plaques. We observed that compression globally increased the lifetime of CCSs as well as the occurrence of static (lifetime >300s) CCSs (Fig. 1a and b), thus confirming previous reports (Ferguson et al., 2017). Because integrin αvβ5, which is necessary for clathrin-coated plaque assembly, could possibly play a role in pressure-induced global loss of CCSs dynamics, we treated cells with Cilengitide, a potent αvβ5 inhibitor. While CCSs were mostly dynamic in Cilengitide-treated cells before pressure, confinement under the agarose gel dramatically increased the lifetime of CCSs as well as the occurrence of stalled CCSs (Fig. 1a and c). These results indicate that CCSs increased lifetime/stabilization under compression is independent of αvβ5 integrin and is most likely the consequence of increased membrane tension. Membrane tension was recently shown to regulate the dynamics of CCSs components in yeast (Aghamohammadzadeh and Ayscough, 2009; Hassinger et al., 2017). To investigate whether pressure also impacts on the dynamics of major CCSs building blocks in our system, we performed fluorescence recovery after photobleaching (FRAP) experiments in cells expressing GFP-tagged μ2-adaptin. We chose to FRAP individual CCSs corresponding to clathrin-coated plaques because the long-lived nature of these structures allows to monitor fluorescence recovery over minutes. In control conditions, fluorescence recovery was fast (half-time recovery, t1/2 ≈ 8s) with a plateau reaching approximately 80%, thus showing that only ~20% of AP-2 complexes were immobile at CCSs (Fig. 1d and e). However, the mobile fraction only reached approximately 60% and half-time recovery was delayed when pressure was applied on cells (t1/2 ≈ 15s; Fig. 1d and e). These results show that cell compression slows down AP-2 turnover in a similar manner as increased membrane tension (Saleem et al., 2015).

**Figure 1.**
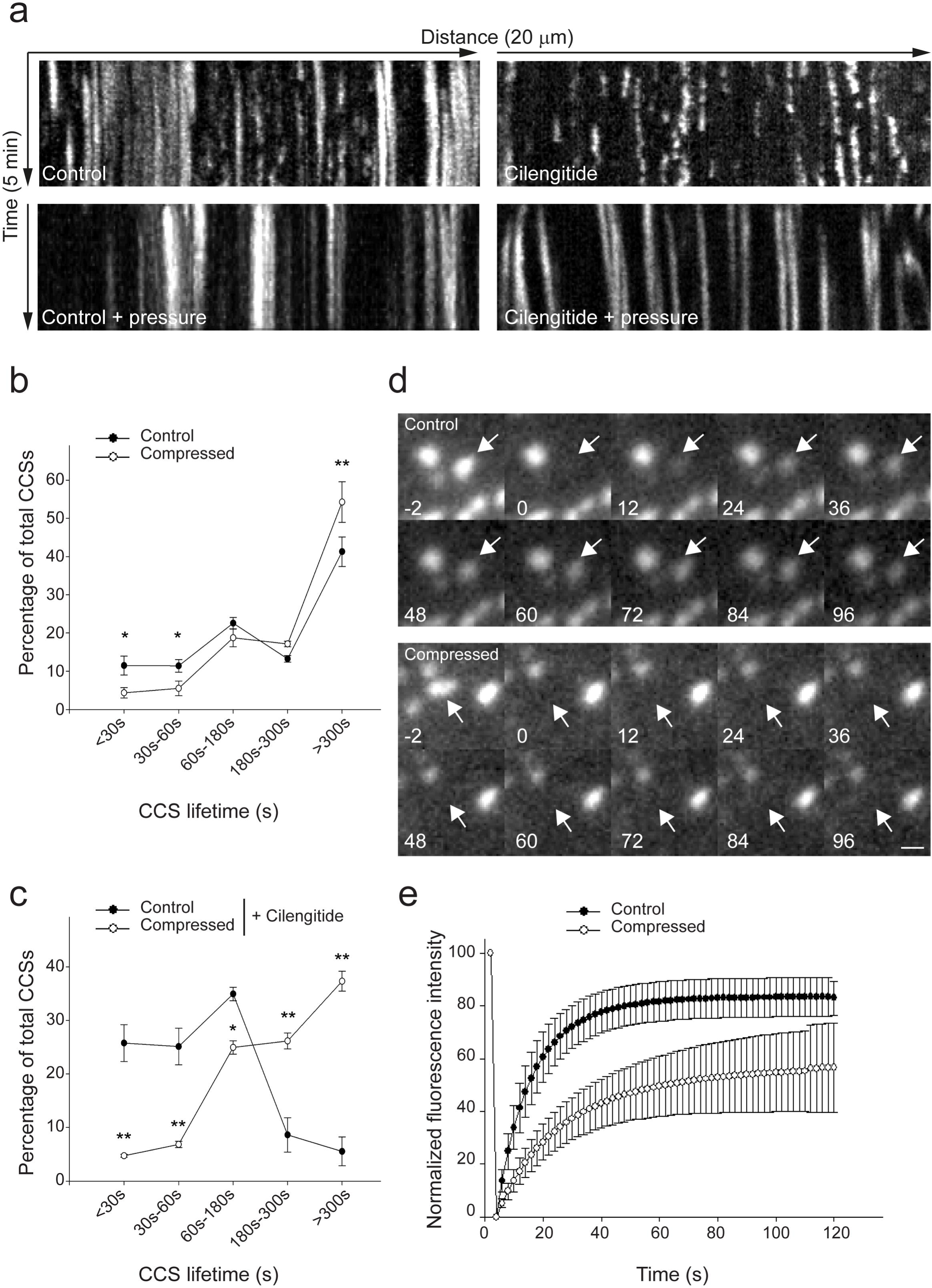
Cell compression reduces CCSs dynamics. **a**, Kymographs showing CCS dynamics in genome-edited HeLa cells expressing endogenous mCherry-tagged μ2-adaptin compressed or not under an agarose plug and treated or not with Cilengitide, as indicated, and imaged by spinning disk microscopy every 5s for 5 min. **b**, **c**, Quantification of the dynamics of CCSs observed as in a (* P<0.05; ** P<0.001, One Way Analysis of Variance – ANOVA. N=3). **d**, Gallery depicting fluorescence recovery of a single mCherry-tagged μ2-adaptin CCS (arrows) after photobleaching in control (upper panels) or compressed (lower panels) cells. Time before or after photobleaching is indicated in seconds. Scale bar: 1μm. **e**, Quantification of fluorescence recovery of mCherry-tagged μ2-adaptin as in d in control or compressed cells as indicated. All results are expressed as mean ± SD.

### Compression leads to CCSs-dependent EGFR signaling

Compressive forces have been reported to activate EGFR and downstream ERK signaling (Tschumperlin et al., 2002; Tschumperlin et al., 2004). Indeed, we observed that compression triggered transient ERK activation (Fig. 2a and b). This held true in several transformed cell lines originated from different tumors, suggesting that compression-induced activation of EGFR might be a widespread characteristic of most cell types (figure S2a and b). We also observed that GFP-tagged ERK transiently translocated from the cytoplasm to the nucleus when HeLa cells were confined under the agarose plug thus confirming the activated status of ERK in these conditions (Fig. S2c and d). However, the mechanoresponsive transcription regulator Yes-activated protein (YAP) remained excluded from the nucleus under pressure suggesting that this pathway is not activated in these conditions (Fig. S2e and f). Pressure-induced ERK activation was dependent on EGFR expression (Fig. 2c and d) as well as on EGFR kinase domain activity as Gefitinib treatment inhibited ERK phosphorylation under compression (Fig. 2e and f).

**Figure 2.**
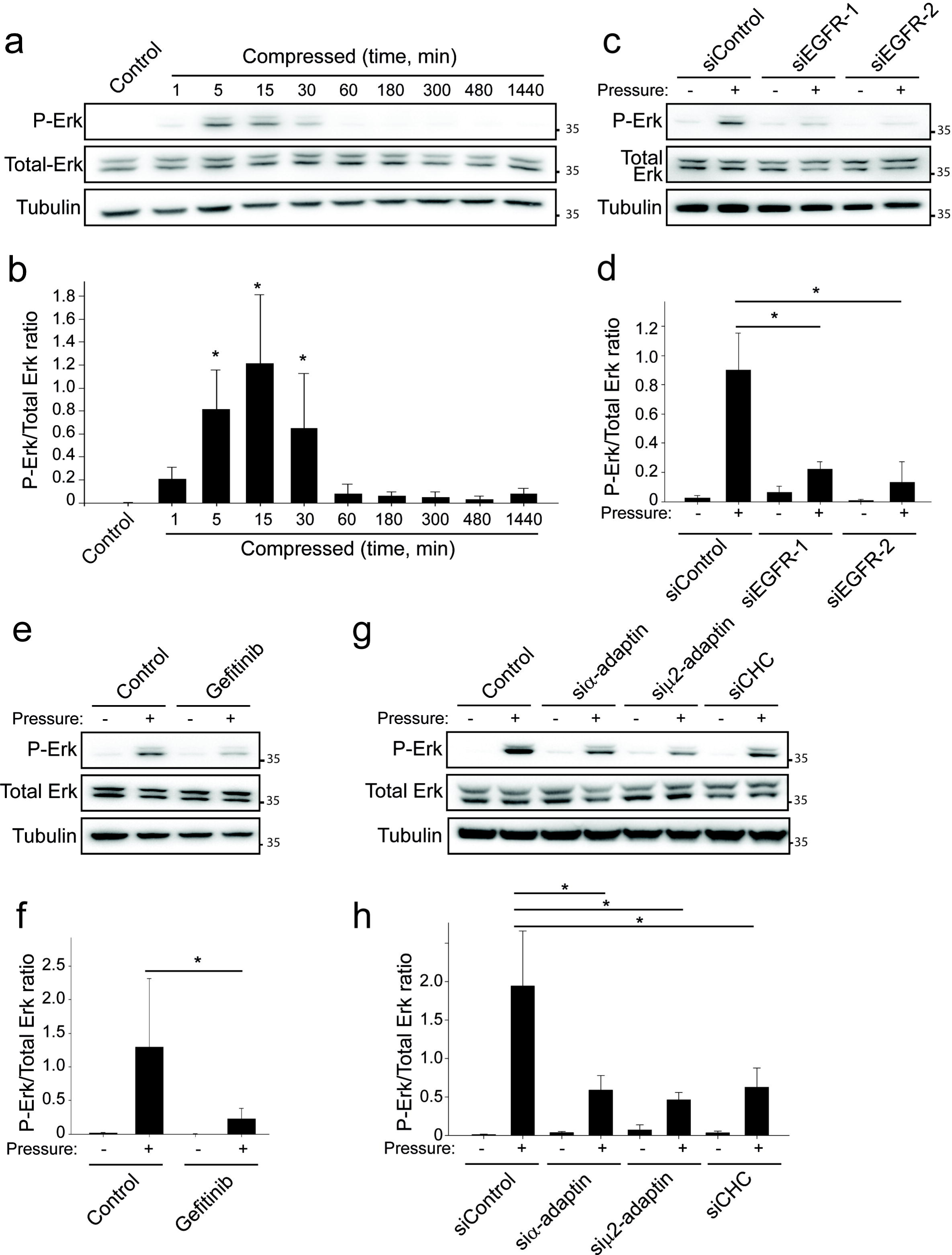
CCSs are required for EGFR-dependent signaling. **a**, Western-blot analysis of phospho-ERK (P-ERK) levels in HeLa cells uncompressed (control) or compressed for different time periods as indicated (representative image of four independent experiments). Total-ERK and tubulin were used as loading controls. **b**, Densitometry analysis of bands obtained in Western-blots as in a. Results are expressed as mean ratio of P-ERK/total ERK ± SD from four independent experiments. **c**, Western-blot analysis of phospho-ERK (P-ERK) levels in HeLa cells compressed or not for 30 min and treated or not with EGFR specific siRNAs as indicated (representative image of four independent experiments). Total-ERK and tubulin were used as loading controls. **d**, Densitometry analysis of bands obtained in Western-blots as in c. Results are expressed as mean ratio of P-ERK/total ERK ± SD from four independent experiments (* *P*<0.05, One Way Analysis of Variance – ANOVA). **e**, Western-blot analysis of phospho-ERK (P-ERK) levels in HeLa cells compressed or not for 30 min and treated or not with Gefitinib for 1 h prior to compression (representative image of four independent experiments). Total-ERK and tubulin were used as loading controls. **f**, Densitometry analysis of bands obtained in Western-blots as in e. Results are expressed as mean ratio of P-ERK/total ERK ± SD from four independent experiments (* *P*<0.05, One Way Analysis of Variance – ANOVA). **g**, Western-blot analysis of phospho-ERK (P-ERK) levels in HeLa cells compressed or not for 30 min and treated or not with AP-2 subunits- or CHC-specific siRNAs as indicated (representative image of four independent experiments). Total-ERK and tubulin were used as loading controls. **h**, Densitometry analysis of bands obtained in Western-blots as in g. Results are expressed as mean ratio of P-ERK/total ERK ± SD from four independent experiments (* P<0.05, One Way Analysis of Variance – ANOVA).

CCSs have been shown to act as platforms that potentiate receptor-mediated signaling, particularly in the case of the EGFR (Sigismund et al., 2008; Vieira et al., 1996). Modulation of CCSs lifetime is an important regulator of receptor signaling output and for instance, long-lived clathrin-coated plaques are more potent than dynamic clathrin-coated pits in supporting signaling (Baschieri et al., 2018). Because compression stalls CCSs dynamics, we reasoned that compression-induced ERK activation could be at least partially due to the increased lifetime of CCSs induced by mechanical stress. Indeed, we observed that AP-2 subunits or clathrin heavy chain (CHC) knockdown reduced ERK activation under compression (Fig. 2g and h and Fig. S2g). Thus, CCSs are required for full EGFR signaling in compressed cells.

### Mechanisms of EGFR recruitment at frustrated CCSs under pressure

EGFR is recruited to CCSs upon ligand-induced activation (Rappoport and Simon, 2009). To ascertain whether compressive stress would lead to the same outcome, genome edited HeLa cells expressing mCherry-tagged, endogenous μ2-adaptin and overexpressing GFP-tagged EGFR were compressed under an agarose plug and monitored using total internal reflection fluorescence (TIRF) microscopy. EGFR quickly accumulated at CCSs upon compression in the absence of serum (Fig. 3a and b, Video 1). Of note, despite the lack of serum, cellular compression still induced ERK activation (Fig. S3a and b). It has been reported that compression induces ectodomain shedding of the EGF-family ligand heparin-binding EGF (HB-EGF), thus leading to autocrine EGFR stimulation (Tschumperlin et al., 2004). HB-EGF shedding is regulated by matrix metalloproteases whose inhibition was shown to prevent EGFR activation following compressive stress (Tschumperlin et al., 2004). Indeed, we observed that Batimastat, a potent and large spectrum inhibitor of matrix metalloproteases, strongly reduced both EGFR recruitment at CCSs (Fig. 3b and c, Video 2) and ERK activation (Fig. 3d and e) upon compression. However, Batimastat did not inhibit ERK activation when EGFR was stimulated using externally added EGF (Fig. S3c and d). Surprisingly, Gefitinib did not prevent the compression-induced EGFR accumulation at CCSs (Fig. 3b and f, Video 2). These latter observations are in favor of a previously proposed model whereby ligand-induced EGFR dimerization is required to interact with the CCS machinery, in a kinase domain activity-independent manner (Wang et al., 2005).

**Figure 3.**
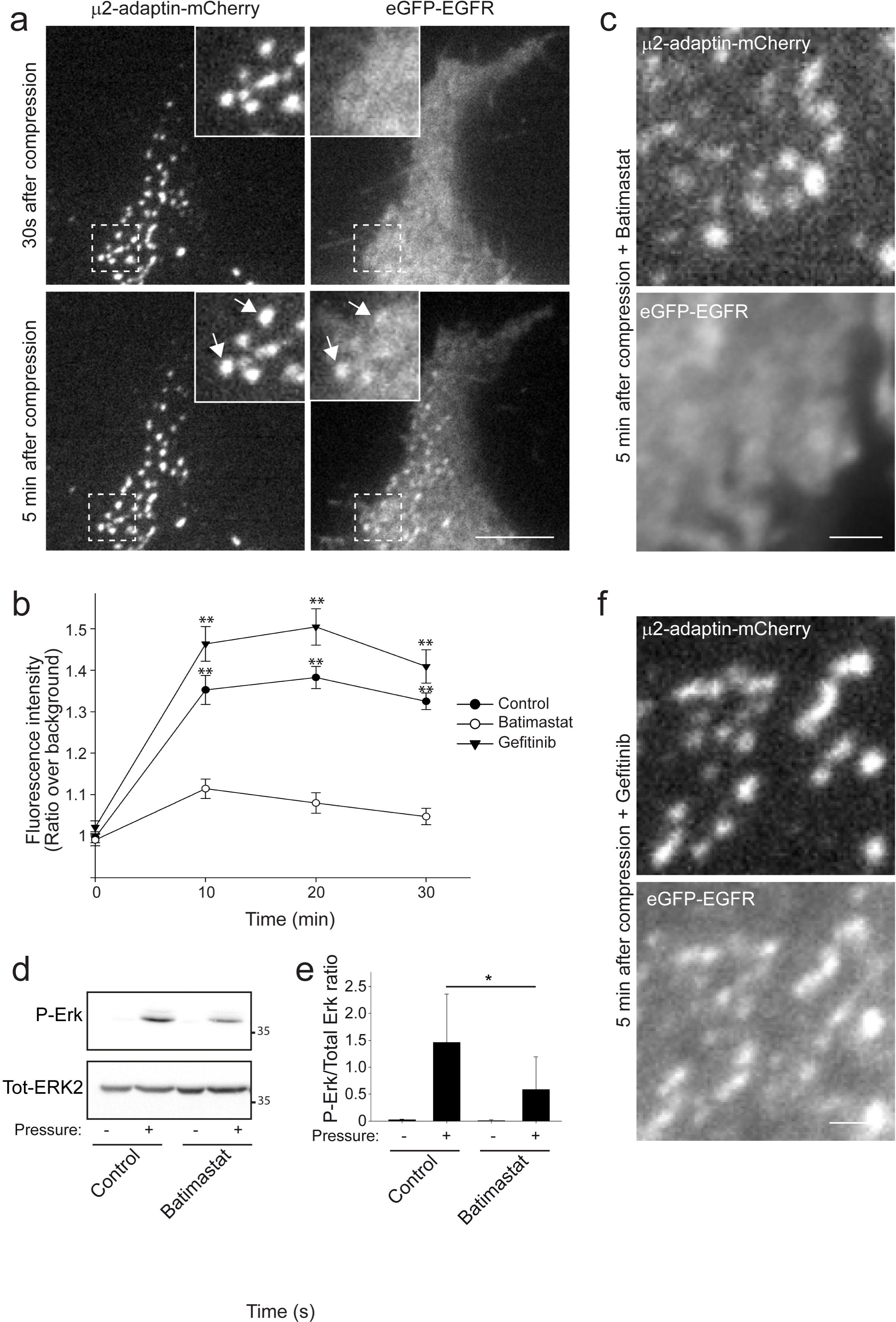
EGFR is recruited at CCSs under compression. **a**, Genome-edited HeLa cells expressing endogenous mCherry-tagged μ2-adaptin were transfected with a plasmid encoding eGFP-tagged EGFR, seeded on glass, compressed under an agarose plug and imaged by TIRF microscopy every 15s for 30 min. Time after compression is indicated. Higher magnifications of boxed regions are shown. Arrows point to EGFR positive CCSs. Scale bar: 8 μm. **b**, Quantification of eGFP-EGFR enrichment at CCSs at the indicated time points after compression in control cells or in cells treated with Batimastat or with Gefitinib, as indicated (** P<0.001 compared to Batimastat, Kruskal-Wallis One Way analysis of variance on ranks. N=3; 20 structures per experiment were analysed). **c**, Genome-edited HeLa cells expressing endogenous mCherry-tagged μ2-adaptin were transfected with a plasmid encoding eGFP-tagged EGFR and seeded on glass. 24 h later, cells were serum starved for at least 2 h, treated with Batimastat for 1 h, then compressed under an agarose plug and imaged by TIRF microscopy every 15s for 30 min. Time after compression is indicated. Scale bar: 1 μm. **d**, Western-blot analysis of phospho-ERK (P-ERK) levels in HeLa cells serum starved for 2 hours, then treated with 10 µM Batimastat for 1h, and subjected or not to compression, as indicated (representative image of four independent experiments). Total-ERK2 was used as a loading control. **e**, Densitometry analysis of bands obtained in Western-blots as in d. Results are expressed as mean ratio of P-ERK/total ERK ± SD from four independent experiments (* P<0.05, One Way Analysis of Variance – ANOVA). **f**, Genome-edited HeLa cells expressing endogenous mCherry-tagged μ2-adaptin were transfected with a plasmid encoding eGFP-tagged EGFR and seeded on glass. 24 h later, cells were serum starved for at least 2 h, treated with 10 µM Gefitinib for 1 h, then compressed under an agarose plug and imaged by TIRF microscopy every 15s for 30 min. Time after compression is indicated. Scale bar: 1 μm.

### Compression-induced CCSs frustration modulates receptor sorting

Frustration of the endocytic machinery could potentially impact any CCS cargo. We therefore aimed at determining whether other receptors could also be recruited at CCSs upon cell compression. We first analyzed the dynamics of the prototypical CCS cargo transferrin receptor (TfR) that is usually constitutively recruited at CCSs in order to be internalized. While mCherry-tagged TfR strongly accumulated at CCSs in control cells, it was excluded from CCSs in compressed cells (Fig. S4a and b). This surprising results suggest that compression modulates receptor sorting at CCSs. Along this line, we observed in FRAP experiments that fluorescence recovery of EGFR-GFP at CCSs was reduced in cells experiencing compression as compared to uncompressed cells stimulated with 10 ng/ml EGF (Fig. S4c). Thus, compression impacts on both CCS component dynamics (Fig. 1d) and CCS cargo dynamics. We next looked at different receptors whose endocytosis is normally triggered by their ligands. G-protein coupled receptor (GPCR) Angiotensin receptor-1 (AT1R), GPCR clathrin adaptor β-arrestin 2, as well as hepatocyte growth factor receptor (HGFR) which are all known to be recruited at CCSs upon stimulation (Eichel et al., 2016; Petrelli et al., 2002) did not accumulate at CCSs under pressure (Fig. 4a-c). These results indicate that compression does not result in the fast activation of these receptors in the absence of their specific ligands.

**Figure 4.**
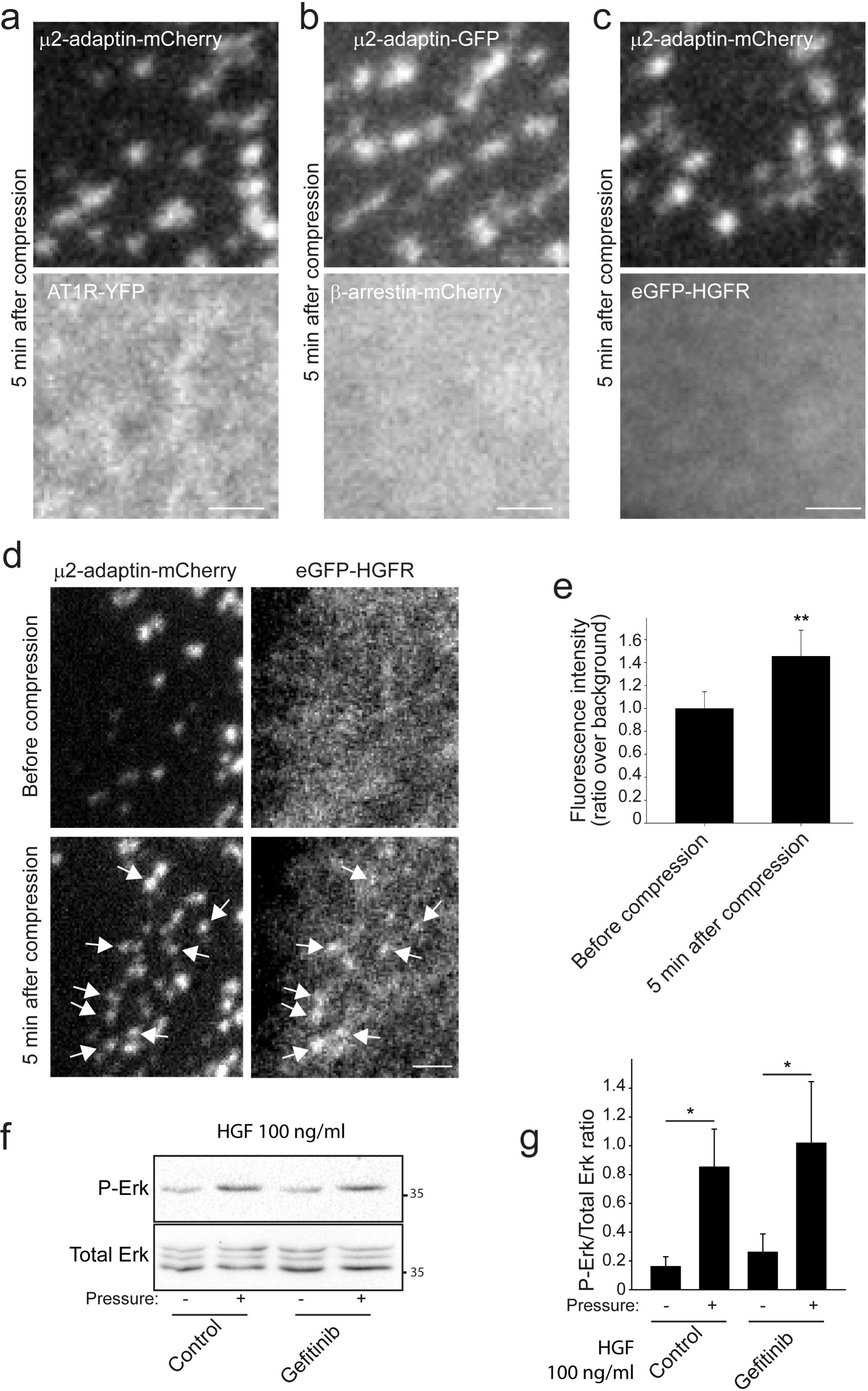
CCSs under compression can serve as signaling platform for different receptors. **a-c**, Genome-edited HeLa cells expressing endogenous mCherry- or GFP-tagged μ2-adaptin were transfected with plasmids encoding YFP-tagged AT1R, mCherry-β-arrestin-2 or GFP-HGFR, as indicated, seeded on glass, compressed under an agarose plug and imaged by TIRF microscopy every 15s for 30 min. Scale bar: 1.5 μm. **d**, Genome-edited HeLa cells expressing endogenous mCherry-tagged μ2-adaptin were transfected with a plasmid encoding eGFP-tagged HGFR. Cells were seeded on glass and 100 ng/ml HGF was added in the culture medium 1h before cells were compressed under an agarose plug and imaged by TIRF microscopy every 15s for 30 min. Time after compression is indicated. Arrows point to HGFR positive CCSs. Scale bar: 2 μm. **e**, Quantification of eGFP-HGFR enrichment at CCSs before or 5 min after compression in control cells treated as in d (** P<0.005, two tailed Student’s T-test. N=3; 20 structures per experiment were analysed.). **f**, Western-blot analysis of phospho-ERK (P-ERK) levels in HeLa cells that were incubated in the presence of HGF for 1h before to be compressed or not and treated or not with Gefitinib, as indicated (representative image of three independent experiments). Total ERK was used as a loading control. **g**, Densitometry analysis of bands obtained in Western-blots as in f. Results are expressed as mean ratio of P-ERK/total ERK ± SD from three independent experiments (* *P*<0.05, One Way Analysis of Variance – ANOVA, Student-Newman-Keuls).

### Compression-induced CCSs frustration controls signaling

We reasoned that, beside the specific case of EGFR, frustrated CCSs could have an impact on signaling downstream of other receptors, provided that receptors’ ligands are present in the system. To test this hypothesis, we incubated HeLa cells in HGF-supplemented medium. While GFP-tagged HGFR did not obviously accumulate at CCSs in these non-acute stimulation conditions, compression led to efficient recruitment of the receptor at CCSs (Fig. 4d and e). HGFR clustering most likely results from low levels of HGFR activation, leading to its progressive accumulation at compression-induced frustrated CCSs that. Importantly, in these conditions, inhibition of EGFR by Gefitinib treatment was not sufficient to block the increased ERK phosphorylation resulting from compression (Fig. 4f and g). This demonstrates that, provided the appropriate ligand is present, receptors other than EGFR can be trapped in compression-induced frustrated CCSs, thus leading to sustained signaling in the ERK pathway.

## DISCUSSION

Here, we confirmed previous findings showing that cell compression leads to frustrated endocytosis, with an accumulation of long-lived CCSs (Ferguson et al., 2017). Several pieces of evidence point to a predominant role of integrins in CCSs frustration through local anchoring of the CCSs machinery to the substrate (Baschieri et al., 2018; De Deyne et al., 1998; Elkhatib et al., 2017). HeLa cells display numerous frustrated CCSs, also termed clathrin-coated plaques, whose formation depends on local enrichment of the αvβ5 integrin (Baschieri et al., 2018). Yet, inhibiting this integrin did not prevent the accumulation of long-lived CCSs in cells experiencing compression. Cell compression most likely results in a dramatic increase in membrane tension that is known to impede CCSs budding (Boulant et al., 2011; Raucher and Sheetz, 1999). Thus, our data strongly suggest that CCSs frustration, as detected in compressed cells, results from increased membrane tension. We also reported that AP-2 dynamics perturbed at frustrated CCSs under compression. This may also result from increased membrane tension as this feature is known to modulate CCS components interaction with the plasma membrane (Saleem et al., 2015). Altered dynamics of CCS components is likely to perturb cargo recruitment at CCSs and, indeed, we observed that the TfR is excluded from CCSs under compression. It is not clear why some receptors like the EGFR and the HGFR can still be recruited at compression-induced frustrated CCSs upon ligand binding, while the TfR, which is normally constitutively addressed to these structures becomes excluded. This may depend on the different types of endocytosis motifs present on receptor cytosolic tails that engage different recognition sites on the AP-2 complex and/or on other CCSs components (Bonifacino and Traub, 2003). In addition, it remains to be elucidated why the dynamics of EGFR is reduced at compression-induced frustrated CCSs. It is possible that the reduced AP-2 dynamics we observed at frustrated CCSs modulates interactions between cargos and the clathrin coat. Along this line, it was recently shown that clathrin dynamic exchange at CCSs is critical to regulate cargo sorting (Chen et al., 2019).

It was previously reported that EGFR becomes activated in bronchial epithelial cells subjected to mechanical stress, leading to strong ERK activation. This activation was shown to be dependent on metalloprotease-regulated HB-EGF shedding leading to paracrine activation of the receptor (Tschumperlin et al., 2004). Our data demonstrate that the same mechanism exist in different transformed cell lines suggesting that compression-induced EGFR activation could play an important role in physiophatological situations like solid tumors that are often characterized by high mechanical pressure. Surprisingly, we observed that EGFR recruitment at CCSs does not depend on the activity of the kinase domain of the receptor. It has long been believed that EGFR autophosphorylation is required for both signaling output and endocytosis of the receptor (Lamaze and Schmid, 1995). Yet, some recent studies have suggested that ligand-induced EGFR dimerization is sufficient to induce the accumulation of the receptor at CCSs, without the need for autophosphorylation of the cytosolic tail (Wang et al., 2002; Wang et al., 2005). Our data clearly support this model.

We additionally showed that CCSs are required for full ERK activation downstream of the EGFR in compressed cells. These observations are in good agreement with previous reports demonstrating that CCSs can serve as signaling platform for EGFR (Sigismund et al., 2008; Vieira et al., 1996). Yet, we previously demonstrated that clathrin-coated plaques can also serve as signaling platform for other receptors (Baschieri et al., 2018). Here, we report that compression-induced frustrated CCSs can potentiate HGFR signaling when HGF is present in the extracellular environment, leading to strong ERK activation even when EGFR kinase activity is inhibited. It is likely that frustrated CCSs trap the few activated HGFRs that, instead of being internalized, progressively accumulate in these stalled structures. This is of special interest in the case of cancer, where cells are often subjected to mechanical stress in an environment that is rich in several growth factors (Shojaei et al., 2010; Straussman et al., 2012; Zhang et al., 2010).

To conclude, we propose that, in complex microenvironments, mechanical stress leads to the activation of the ERK signaling pathway not only because of HB-EGF shedding and paracrine activation of the EGFR, but also because compression-induced CCSs can trap and potentiate signaling by many other receptors.

## METHODS

### Cell lines and constructs

HeLa cells (a gift from P. Chavrier, Institut Curie, Paris, France; ATCC CCL-2), genome-edited HeLa cells engineered to expressed an endogenous GFP-tagged or mCherry-tagged µ2 subunit, HepG2 cells (ATCC HB-8065), Caco-2 cells (ATCC HTB-37), Hs578t cells (a gift from C. Lamaze, Institut Curie, Paris, France), were grown in DMEM Glutamax supplemented with 10% foetal calf serum at 37°C in 5% CO2. For microscopy, cells were serum-starved for at least 2h before the experiment. All cell lines have been tested for mycoplasma contaminations. mCherry-TfR was a gift from Michael Davidson (Addgene plasmid #55144). GFP-ERK2 was a gift from Dr.Hesso Farhan. EGFR-GFP was a gift from Alexander Sorkin (Addgene plasmid # 32751). pLenti-MetGFP was a gift from David Rimm (Addgene plasmid # 37560). pEGFP-C3-YAP2 was a gift from Marius Sudol (Addgene plasmid # 19055). AT1R-YFP and β-arrestin2-mCherry were a kind gift of Dr. Mark Scott (Institut Cochin, Paris, France).

Plasmids were transfected 24h after cell plating using either Lipofectamine 3000 according to the manufacturer’s instructions or electroporating cells in suspension using AMAXA nucleofector Kit V according to the manufacturer’s instructions. Alternatively, linear Polyethylenimine (PEI - MW 25.000 – Polysciences Cat. Nr. 23966) at 1 mg/ml was used to transfect 50 % confluent cells in a 6 well plate according to the following protocol: 2 µg of DNA were added to 100 µl of OptiMEM, followed by addition of 4 µl of PEI, vortex and incubation for 10 minutes at RT prior to add the mix to the cells.

### Antibodies and drugs

Rabbit polyclonal antibodies anti tot-ERK1/2 (Cat. Nr. 9102) and P-ERK1/2 (Cat. Nr. 9101) were purchased from Cell Signalling. Mouse monoclonal anti tot-ERK1/2 (Cat. Nr. 13-6200) was purchased from Thermo Fisher. Mouse monoclonal anti-α-Tubulin (Cat. Nr. T9026) was purchased from Sigma. Mouse monoclonal anti-clathrin heavy chain (CHC, Cat. Nr. 610500) was purchased from BD Biosciences. Mouse monoclonal anti-α-adaptin (Cat. Nr. ab2807) and rabbit monoclonal anti-EGFR (Cat. Nr. ab2430) were purchased from Abcam. Gefitinib (Cat. Nr. CDS022106) was purchased from Sigma and used at a final concentration of 10 µM. Cilengitide was purchased from Selleckchem (Cat. Nr. S7077) and used at a final concentration of 10 µM. Human recombinant HGF (Cat. Nr. 1404) was purchased from Sigma and used at a final concentration of 100 ng/ml. For HGF experiments, cells were previously serum-starved for at least 2h and HGF was added to the serum-free medium for 1h before experiment. Sir-DNA (Spirochrome sc-007) was used at 0.5 µM to stain nuclei for live cell imaging. Cells were incubated for 20 minutes in the presence of Sir-DNA, then washed once with PBS to eliminate excess Sir-DNA and put back in DMEM.

### In vitro compression experiments

To investigate the effect of compressive stress on cell behavior, an under-agarose assay was used (Heit and Kubes, 2003). Cells were plated either in 6-well cell culture plates or in glass-bottom dishes (µ-Dish Cat Nr 190301, Ibidi). 24h hours later, cells were subjected to mechanical stress by using an agarose plug overlaid with the weight necessary to reach a pressure of approximately 1000 Pa (agarose plug of 1.9 cm², corresponding to the area of one well of a 24-well, and weight of 20 g). To prepare agarose gels, agar was weighted and dissolved in DMEM Glutamax to a final concentration of 2.4%. The mixture was then cast in an empty dish or well and cooled at room temperature. Agar disks were sterilized under UV light and equilibrated at 37°C before use. For western blots, cells were subjected to compression for 30 minutes prior to cell lysis, unless otherwise stated. For video microscopy, videos were started 30 sec before applying the compressive stress. Compressed cells were then imaged for 30 min acquiring one image every 15 sec. Alternatively, for CCSs dynamics and for FRAP experiments, videos were acquired before and under compression and the videos before compression were compared to the videos under compression. For ERK2 and YAP nuclear enrichment, cells were treated with Sir-DNA as described above and imaged by spinning disk microscopy. Cells were manually segmented and the Sir-DNA channel was used for the segmentation of the nucleus. Cytoplasm segmentation was obtained by subtracting the nucleus area from the cell area. ERK or YAP fluorescence was measured over time in the nucleus and in the cytoplasm, then the nuclear enrichment (nuclear/cytosolic signal ratio) was calculated for each cell.

### Western Blots

For Western Blot experiments, cells were lysed in ice cold MAPK buffer (100mM NaCl, 10 nM EDTA, 1% IGEPAL ® CA-630, 0.1% SDS, 50mM TRIS-HCl pH 7.4) supplemented with protease and phosphatase inhibitors. Protein concentration was measured with Pierce™ Coomassie Plus (Bradford) Assay Kit (Cat Nr 1856210) according to the manufacturer’s instructions in order to load equal amount of proteins. Antibodies were diluted at 1:1000 in PBS - 0.1% Tween - 5% BSA or 5% non-fat dried milk. For stripping, membranes were incubated in a commercial stripping buffer (Cat. Nr ST010; Gene Bio-Application) according to the manufacturer’s instructions. Western-blot quantifications were done in FIJI.

### RNA interference

For siRNA depletion, 200 000 cells were plated in 6 well plates. After 24 h, cells were treated with the indicated siRNA (30 nM) using RNAimax (Invitrogen, Carlsbad, CA) according to the manufacturer’s instruction. Protein depletion was maximal after 72 h of siRNA treatment as shown by immunoblotting analysis with specific antibodies. To deplete CHC, α-adaptin or µ2-adaptin, cells were transfected once as described above and then a second time, 48 hours later, with the same siRNAs. In this case, cells were analyzed 96 hours after the first transfection. The following siRNAs were used: µ2-adaptin, 5’-AAGUGGAUGCCUUUCGGGUCA-3’; Clathrin heavy chain (CHC), 5’GCUGGGAAAACUCUUCAGATT-3’; α-adaptin, 5’-AUGGCGGUGGUGUCGGCUCTT-3’; Epidermal growth factor receptor (EGFR) 5’-GAGGAAAUAUGUACUACGA-3’ (EGFR-1) and 5’-GCAAAGUGUGUAACGGAAUAGGUAU-3’ (EGFR-2); non-targeting siRNAs (siControl), ON-TARGETplus Non-Targeting SMARTpool siRNAs (Dharmacon D-001810-01).

### Spinning disk microscopy of live cells

For CCSs dynamics, cells were imaged at 5 s intervals for the indicated time using a spinning disk microscope (Andor) based on a CSU-W1 Yokogawa head mounted on the lateral port of an inverted IX-83 Olympus microscope equipped with a 60x 1.35NA UPLSAPO objective lens and a laser combiner system, which included 491 and 561 nm 30 mW DPSS lasers (Andor). Images were acquired with a Zyla sCMOS camera (Andor). Alternatively, cells were imaged on a Nikon Ti2 Eclipse (Nikon France SAS, Champigny sur Marne, France) inverted microscope equipped with a 60x NA 1.40 Oil objective WD 0.130, a sCMOS PRIME 95B camera (Photometrics, AZ, USA) and a dual output laser launch, which included 405, 488, 561 and 642 nm 30 mW lasers. Both microscopes were steered by Metamorph 7 software (MDS Analytical Technologies, Sunnyvale, CA, USA).

For CCS dynamics quantification, the lifetime of CCSs was measured using the TrackMate plugin of ImageJ (Tinevez et al., 2017). Tracks below 5 seconds of duration (detected on only 1 frame) were discarded. Measured individual lifetimes were pooled into two groups: the “dynamic” group corresponding to structures with a lifetime below the duration of the movie (5 min) and the “static” group with a lifetime of 5 min. Of note, the relative percentage of dynamic versus static structures depends on the duration of the movie because static structures are only counted once while dynamic structures continuously nucleate and disappear during the movie. To circumvent this limitation, all quantifications of CCS dynamics represent the relative number of static or dynamic events detectable at the plasma membrane at a given time point. At least 1000 CCSs from at least 5 cells per conditions and per experiments were tracked in 3-5 independent experiments. Data are expressed as mean ± SD.

### Total internal reflection fluorescence microscopy (TIRF) and Fluorescence Recovery After Photobleaching (FRAP)

For total internal reflection fluorescence microscopy (TIRF), HeLa cells transfected with the indicated plasmids were imaged through a 100x 1.49 NA APO TIRF WD 0.13-0.20 oil objective lens on a Nikon Ti2 Eclipse (Nikon France SAS, Champigny sur Marne, France) inverted microscope equipped with a sCMOS PRIME 95B camera (Photometrics, AZ, USA) and a dual output laser launch, which included 405, 488, 561 and 642 nm 30 mW lasers, and driven by Metamorph 7 software (MDS Analytical Technologies, Sunnyvale, CA, USA). A motorized device driven by Metamorph allowed the accurate positioning of the illumination light for evanescent wave excitation.

For TIRF-FRAP experiments, one CCS was manually selected and subjected to 100% laser power (30 mW laser) scan in order to have a bleaching of at least 80% of the fluorescence. One frame was collected before photo-bleaching, and 60 frames were collected after bleaching to analyze fluorescent recovery at the frequency of 1 frame/2 sec. The data were analyzed using the ImageJ FRAP Profiler plugin (McMaster University, Canada) to extract recovery curves and calculate the half-time recovery.

### Statistical analyses

Statistical analyses in Fig.1 (panels b, c), Fig.2 (panels b, d, f, h), Fig.3 (panel e), Fig.4 (panel g), Supplementary Fig.3 (panel d) were performed using One Way Analysis of Variance (ANOVA). Statistical analyses in Fig.3 (panel b) were performed using Kruskal-Wallis One Way analysis of variance on ranks. Statistical analyses in Fig.4 (panel e), Supplementary Fig.2 (panel b) Supplementary Fig.3 (panel b), and Supplementary Fig.4 (panel b) were performed using two tailed Student’s T-test. All data are presented as mean of at least four independent experiments ± SD, except otherwise stated. All statistical analyses were performed using SigmaPlot software.

### Data availability

The authors declare that all data supporting the findings of this study are available within the article and its supplementary information files or from the corresponding author upon reasonable request.

## Acknowledgment

We thank the imaging facilities of Gustave Roussy for help with image acquisition. Core funding for this work was provided by the Gustave Roussy Institute and the Inserm and additional support was provided by grants from ATIP/Avenir Program, la Fondation ARC pour la Recherche sur le cancer, Le Groupement des Entreprises Françaises dans la LUtte contre le Cancer (GEFLUC) and from the Agence Nationale de la Recherche (ANR-15-CE15-0005-03) to GM. This project was supported by grant "Taxe d’apprentissage Gustave Roussy - 2017 - DLD"

F.B and D.L designed and performed experiments, analysed results and wrote the manuscript. N.E performed experiments. G.M supervised the study, designed experiments and wrote the manuscript.

The authors declare no competing interests. Correspondence and requests for materials should be addressed to or to

## Supplementary figure legends

Figure S1. **Analysis of cell compression efficiency. a**, Wide-field image of one HeLa cell compressed under and agarose plug. Arrows points to blebs at the plasma membrane. Scale bar: 5μm. **b**, HeLa cells treated with Sir-DNA were imaged by spinning disk microscopy before or after compression under an agarose plug, as indicated. Arrows point to nuclear blebs. Scale bar: 10μm.

Figure S2. **Analysis of ERK and YAP behavior under pressure. a**, Western-blot analysis of phospho-ERK (P-ERK) levels in the indicated cell lines compressed or not for 30 min (representative image of three independent experiments). Total-ERK was used as loading control. **b**, Densitometry analysis of bands obtained in Western-blots as in a. Results are expressed as mean ratio of P-ERK/total ERK ± SD from 3 independent experiments (* *P*<0.05; ** P<0.001, two-tailed paired t-test). **c**, HeLa cells transfected with a plasmid encoding eGFP-tagged ERK2 were imaged by spinning disk microscopy before (upper panel) or after (lower panel) being compressed under an agarose plug. Scale bar: 10μm. **d**, Quantification of eGFP-ERK2 enrichment in the nucleus at the indicated time points after compression. 22 cells from 3 independent experiments were quantified. Results are expressed as mean ± SE. **e**, HeLa cells transfected with a plasmid encoding eGFP-tagged YAP were imaged by spinning disk microscopy before (upper panel) or after (lower panel) being compressed under an agarose plug. Scale bar: 10μm. **f**, Quantification of eGFP-YAP enrichment in the nucleus at the indicated time points after compression. 35 cells from 3 independent experiments were quantified. Results are expressed as mean ± SE. **g**, HeLa cells were transfected with the indicated siRNAs. 72 h after transfection, cells were lysed and subjected to Western-blot analysis using the indicated antibodies. Tubulin was used as loading control.

Figure S3. **EGFR activation under compression is serum-independent. a**, Western-blot analysis of phospho-ERK (P-ERK) levels in starved HeLa cells compressed or not in FCS-free medium, as indicated (representative image of four independent experiments). Total ERK was used as a loading control. **b**, Densitometry analysis of bands obtained in Western-blots as in a. Results are expressed as mean ratio of P-ERK/total ERK ± SD from four independent experiments (* *P*<0.05, One Way Analysis of Variance – ANOVA). **c**, Western-blot analysis of phospho-ERK (P-ERK) levels in HeLa cells serum starved for at least 2h, treated or not with batimastat for 1 h, and stimulated with 10 ng/ml EGF or not as indicated (representative image of three independent experiments). Total-ERK was used as loading control. **d**, Densitometry analysis of bands obtained in Western-blots as in c. Results are expressed as mean ratio of P-ERK/total ERK ± SD from three independent experiments (** *P*<0.01, One Way Analysis of Variance – ANOVA, Student-Newman-Keuls).

Figure S4. **Alteration of receptor sorting and dynamics under compression. a**, Genome-edited HeLa cells expressing endogenous eGFP-tagged μ2-adaptin were transfected with a plasmid encoding mCherry-tagged TfR, seeded on glass and imaged by TIRF microscopy before (upper panel) or 5 min after (lower panel) compression. Arrows point to TfR positive CCSs. Arrowheads point to TfR-positive, AP-2-negative structures most likely corresponding to endosomes. Scale bar: 2 μm. **b**, Quantification of mCherry-TfR enrichment at CCSs before or 5 min after compression in cells treated as in a (* P<0.005, two tailed Student’s T-test. N=3; 80 to 100 structures per experiment were analysed.). **c**, Quantification of fluorescence recovery after photobleaching of the eGFP-EGFR fluorescence in individual CCSs in cells stimulated with EGF or compressed under an agarose plug, as indicated. All results are expressed as mean ± SD.

## Supplementary Videos

Video 1. **EGFR recruitment to CCSs.** Genome-edited HeLa cells expressing endogenous mCherry-tagged μ2-adaptin were transfected with a plasmid encoding eGFP-tagged EGFR, seeded on glass and starved for 1h. Cells were then compressed under an agarose plug (1 kPa pressure) and imaged by TIRF microscopy every 15s for 30 min. Scale bar: 10 μm.

Video 2. **EGFR recruitment to CCSs in the presence of Gefitinib or Batimastat.** Genome-edited HeLa cells expressing endogenous mCherry-tagged μ2-adaptin were transfected with a plasmid encoding eGFP-tagged EGFR, seeded on glass, starved for 1h in the presence of 10 µM Gefitinib (left) or 10 µM Batimastat (right). Cells were then compressed under an agarose plug (1 kPa pressure) and imaged by TIRF microscopy every 15s for 30 min. Scale bar: 10 μm.

