## Supplementary figures and images for "Endocytosis frustration potentiates compression-induced receptor signaling"

### Supplementary figures

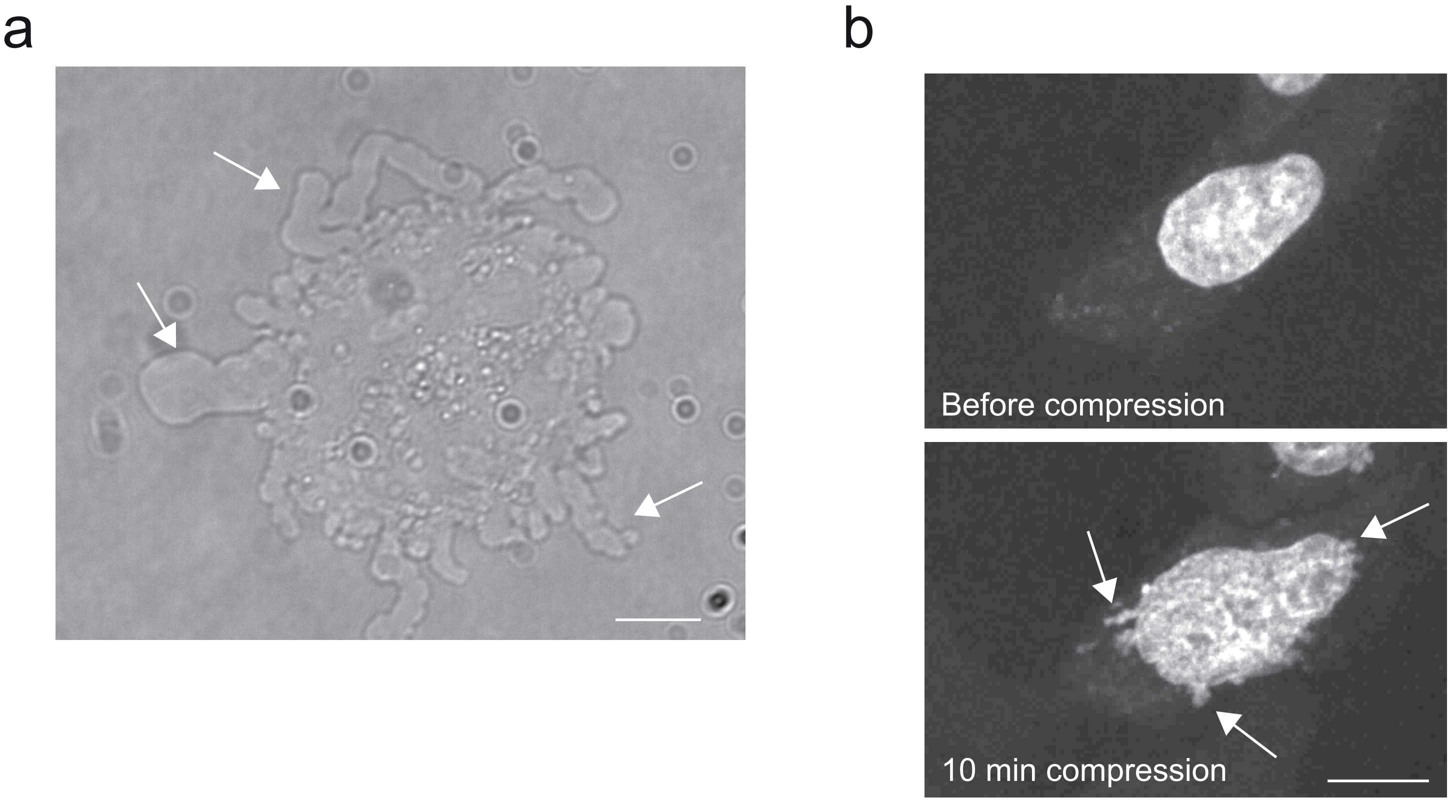


Figure S1

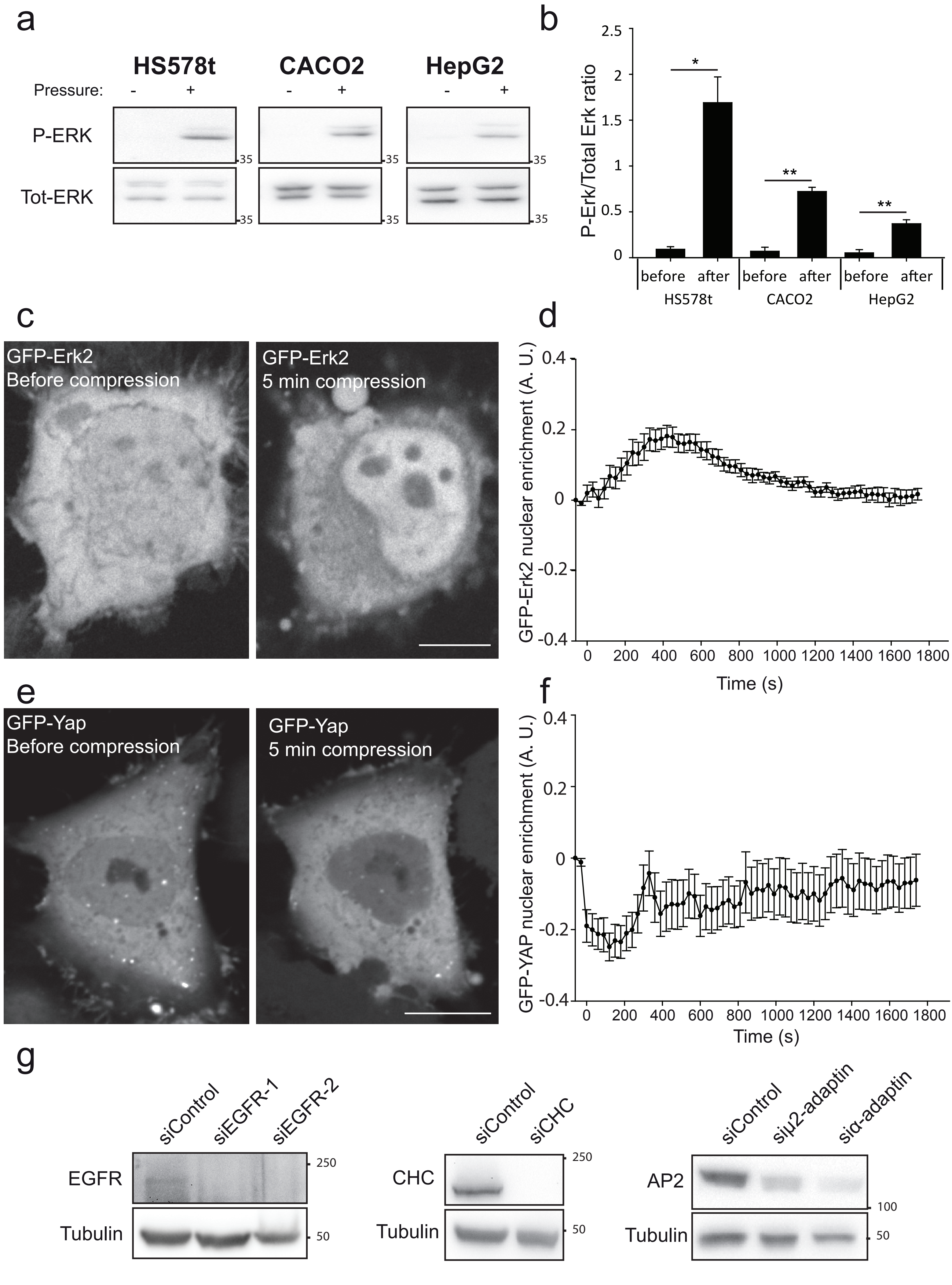

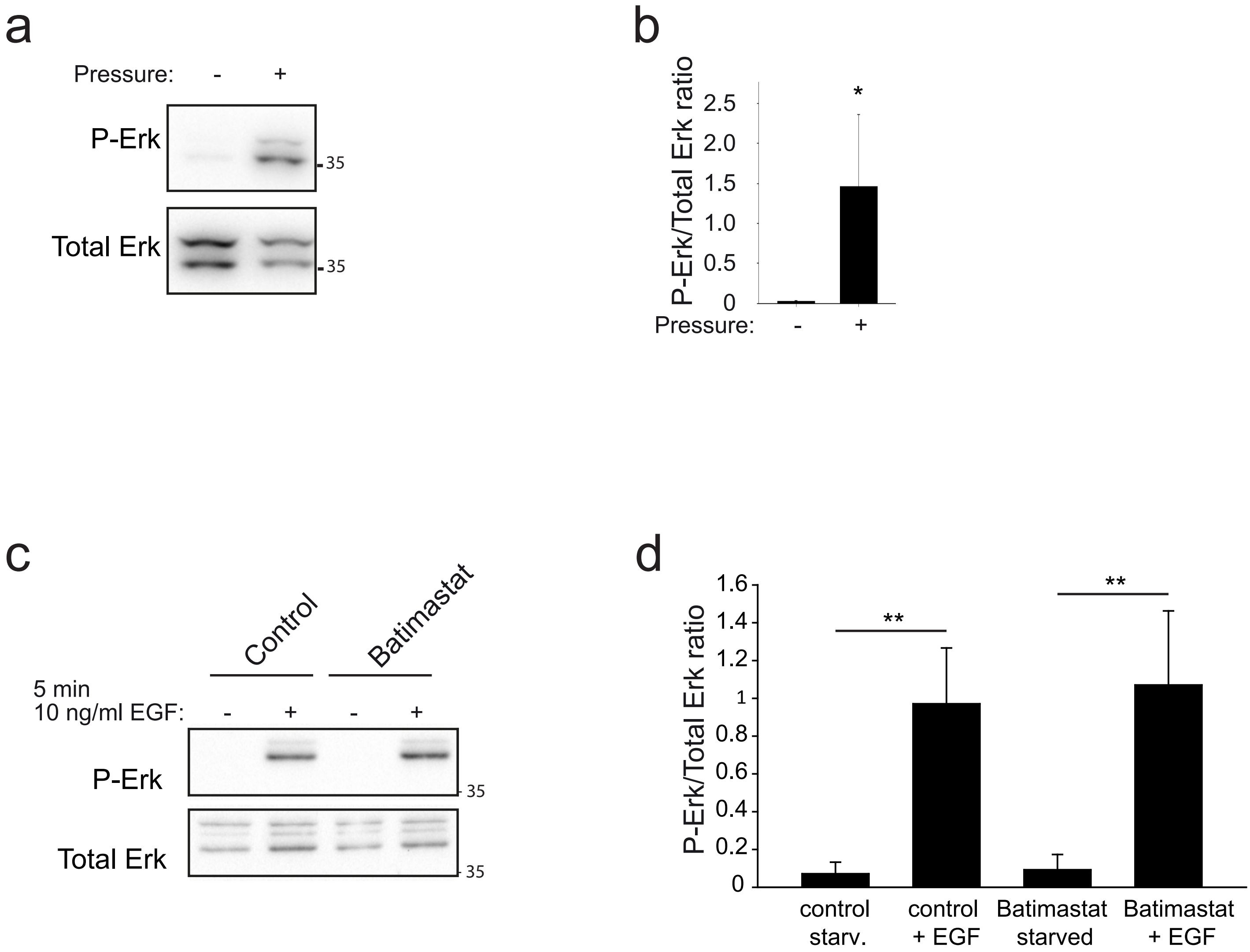

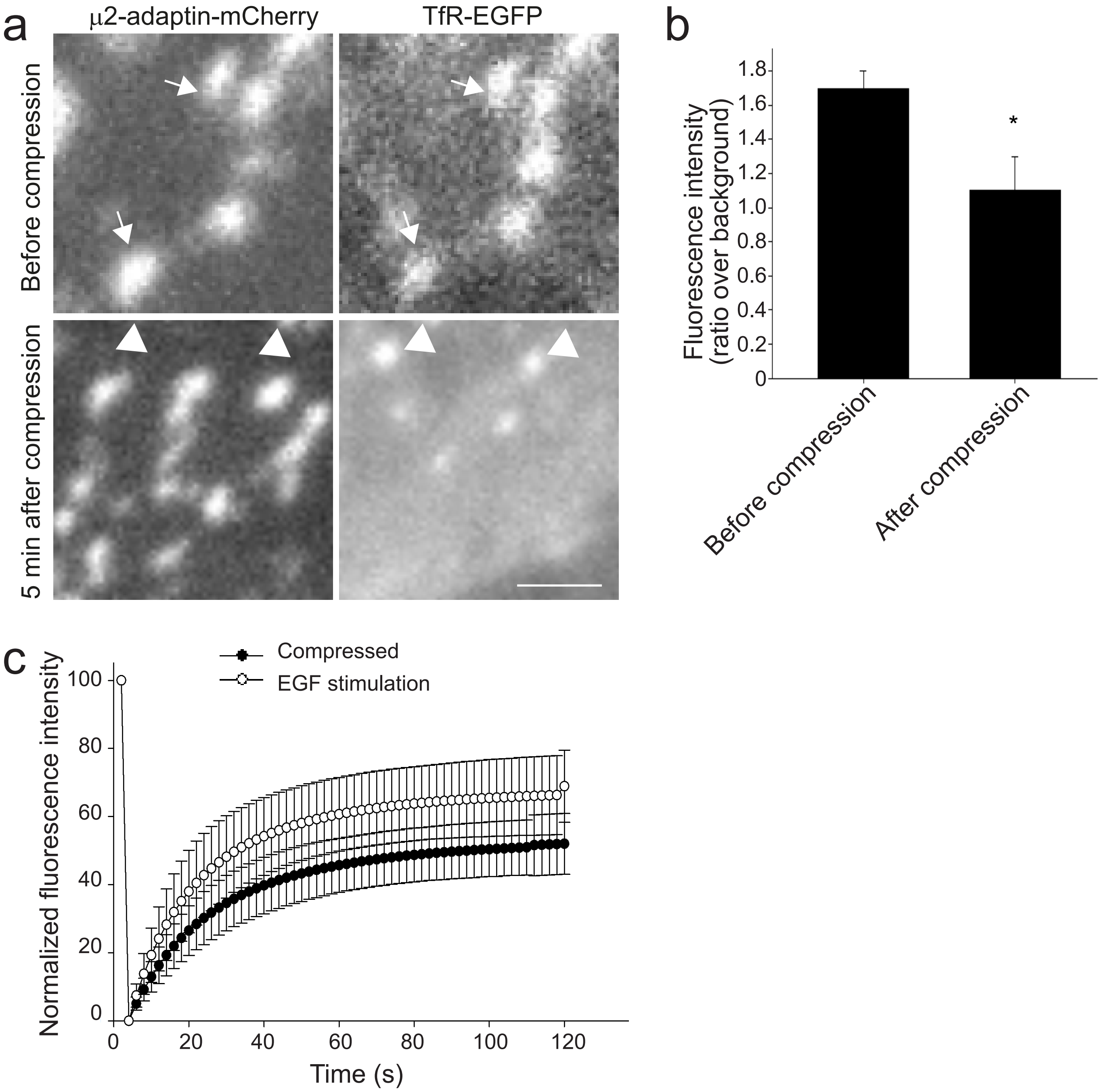
